# Hot, humid air decontamination of aircraft confirmed that high temperature and high humidity are critical for inactivation of infectious, enveloped ribonucleic acid (RNA) virus

**DOI:** 10.1101/2020.07.20.212365

**Authors:** Tony L. Buhr, Alice A. Young, Erica Borgers-Klonkowski, Neil L. Kennihan, Harold K. Barnette, Zachary A. Minter, Matthew D. Bohmke, Emily B. Osborn, Shelia M. Hamilton, Monique B. Kimani, Mark W. Hammon, Charles T. Miller, Ryan S. Mackie, Jennifer M. Innocenti, Misty D. Bensman, Bradford W. Gutting, Samuel D. Lilly, Emlyn E. Hammer, Vanessa L. Yates, Brooke B. Luck

## Abstract

**Aims:** To develop infectious (live/dead) enveloped virus test indicators and Response Surface Methodology (RSM) models that evaluate survival of an enveloped ribonucleic acid (RNA) virus on contaminated aircraft materials after exposure to hot, humid air (HHA).

**Methods and Results:** Enveloped RNA bacteriophage Phi6 (Φ6) was dried on wiring insulation, aircraft performance coating (APC), polypropylene, and nylon at ≥ 8 log_10_ plaque-forming units (PFU) test coupon^-1^. Only 2.4 log_10_ inactivation was measured on APC at 70°Celsius (°C), 5% relative humidity (RH) after 24 h. In contrast, HHA RSM models showed a 90% probability of a 7-log_10_ inactivation at ≥63°C, 90% RH after 1 h, and decontamination kinetics were similar across different materials. HHA decontamination of C-130 and C-17 aircraft showed >7 log_10_ and ≥5.9 log_10_ inactivation of enveloped virus on 100 and 110 test indicators, respectively, with a 1-h treatment, excluding ramp-up and ramp-down times.

**Conclusions:** Enveloped RNA virus test indicators were successfully developed, lab tested for HHA decontamination, analyzed for RSM, and field-tested in aircraft demonstrations.

**Significance and Impact of the Study:** The utility of HHA decontamination was demonstrated after inactivating enveloped RNA virus on aircraft with a 1-h HHA treatment within aircraft temperature and RH limits.

## INTRODUCTION

There is a need to develop aircraft decontamination procedures for enveloped RNA viruses ranging from hemorrhagic fever viruses to coronaviruses (Centers for Disease Control 2019; National Business Aviation Association 2020). A candidate procedure for aircraft decontamination is hot, humid air (HHA), which has been tested for *Bacillus* spore decontamination (Buhr et al. 2012, 2015, 2016). HHA decontamination is highly plausible for enveloped virus decontamination since enveloped viruses are considered much easier to kill than spores (Spaulding 1957). In order to develop HHA decontamination profiles and test in aircraft field demonstrations, a biosafety level 1 (BSL-1) enveloped RNA virus was needed as a test indicator to meet the requirements imposed by biosafety and environmental reviews for field-testing. Field test approval is a lengthy process that can require decades of compiled research to approve BSL-1 organisms for field testing, regardless of whether the organism is contained or released during the test (Bishop and Robinson 2014; Buhr et al. 2016). It is particularly challenging to justify and approve enveloped virus strains for rapid field-testing (2 months for biosafety approval at different facilities, enveloped virus test indicator preparation, and 2 aircraft field tests including results) during a pandemic such as the SARS-CoV-2/COVID19 pandemic. Furthermore, active, infectious virus was needed for a live/dead (infectious/non-infectious for viruses) assay since live/dead assays are the hallmark of decontamination testing.

Φ6 is a BSL-1 enveloped RNA virus originally isolated in a bean field as a lytic virus that infected the plant pathogenic bacterium *Pseudomonas syringae* pathovar *phaseolicola* (Vidaver et al. 1973; Van Etten et al. 1976; Mindich 2004). Early hopes to produce the virus in large quantities to spray on bean fields as an environmentally friendly biocontrol agent were never commercially realized. However, due to its rare combination as a BSL-1, enveloped RNA virus it has been proposed as a general surrogate for a number of different enveloped RNA viruses particularly for field-testing (Gallandat and Lantagne 2017). The Φ6 envelope structure is similar to many other enveloped viruses as the envelope consists of a glycoprotein/protein-embedded lipid membrane and the host cell has similar temperature sensitivity to mammalian cells at around 40°C. This is important since the envelope components are considered a primary target for inactivation by many different decontaminants (McDonnell and Burke 2011; Wiggington et al. 2012). Here Φ6 was prepared to develop BSL-1 enveloped virus test indicators at ≥ 7.6 log_10_ coupon^-1^, test HHA decontamination parameters, develop response surface models for HHA decontamination, and measure enveloped virus inactivation during C-130 and C-17 aircraft field tests.

## MATERIALS AND METHODS

### Φ6 and Host Cell Preparations

Φ6 and its host organism *P. syringae* pathovar *phaseolicola* HB10Y (HB10Y), causal agent of halo blight of the common bean, *Phaseolus vulgaris*, were isolated in Spain. Both were a kind gift from Dr. Leonard Mindich at Rutgers University, New Jersey Medical School. HB10Y was prepared by inoculating 100-200 ml of 3% tryptic soy broth (TSB; Fluka PN#T8907-1KG) in a 1-liter (l) smooth-bottom Erlenmeyer flask with a high efficiency particulate air filter cap. Cultures were incubated at 26±2°C, 200 revolutions (rev) minute (min)^-1^ for 20±2 hours (h). 11.1 ml of 100% glycerol (Sigma PN #G7757-500ML) were added per 100 ml of host culture. Final concentration of glycerol was 10%. One-ml aliquots of HB10Y were pipetted into screw-cap microfuge tubes with O-rings, and stored at −80°C. HB10Y samples were titered prior to freezing by serially diluting samples in 10 millimolar (mM) of 4-(2-hydroxyethyl)-1-piperazineethanesulfonic acid (HEPES, Sigma PN#H4034-100G) + 10% Sucrose (Sigma PN #S7903-250G), pH 7.0, and plating on Tryptic Soy Agar (TSA; Hardy Diagnostics, Santa Maria, CA). Plates were inverted and incubated at 26±2°C for 48±2 h to show titers of ~10^9^ cells ml^-1^. After freezing, tubes were thawed at room temperature (RT, 22±3°C), serially diluted and plated to show sustained viability after long-term storage at −80°C.

Φ6 was prepared after inoculating broth cultures of HB10Y. A frozen stock prep of HB10Y was thawed at 22±3°C. HB10Y was added either directly from a frozen stock or by transferring a single colony from a streaked TSA plate to 100-200 ml of 3% TSB in a 1-l smooth-bottom Erlenmeyer flask with a HEPA cap and incubated at 26±2°C, 200 rev min^-1^ overnight. Cells were then diluted and grown to mid-log-phase. The host flask was inoculated with 0.5-1 ml of Φ6 at a concentration of ~1-2e11 plaque-forming units (PFU) ml^-1^ at the time of inoculation. The culture was incubated at 26±2°C, 200 rev min^-1^ for 24±2 h. The Φ6 preparation was stored at 4°C until after titering was completed. After titer determination was completed, typically around 1-2e11 PFU ml^-1^, then 1-1.3 ml volumes were aliquoted into 1.5-ml screw-cap tubes with O-rings, inverted and stored at −80°C.

### Environmental Test Chamber Setup and Validation

Thermotron SM-8-8200 (Thermotron Industries, Holland, MI, USA) environmental chambers were used to control temperature and RH as described (Buhr et al. 2015).

### Coupon Materials and Sterilization

Square 2 centimeter (cm) x 2 cm coupons of different test materials or the inside surface of 50-ml Techno Plastic Products AG, Switzerland (TPP^®^) polypropylene conical tubes were inoculated with >8 log_10_ Φ6 virus inoculum. The inoculated 2 cm x 2 cm coupons were set inside sterile TPP^®^ tubes during testing (Buhr et al. 2012, 2015, 2016). Materials for inoculation included aluminum 2024-T3 coupons painted with water-based aircraft performance coating (APC), stainless steel 304 coupons painted with Navy Top Coat (NTC) (Coatings Group at the University of Dayton Research Institute (UDRI), Dayton, OH, USA), wiring insulation (Kapton Film Type HN, 1 mil, Reference No. 6197844-00 from Cole Parmer, Vernon Hills, IL, USA), and nylon webbing (nylon) (US Netting, Erie, PA, USA) with the ends of each coupon cauterized to prevent fraying. Prior to inoculating and testing, coupons were rinsed with 18 mega-Ohm-cm, de-ionized water, placed on absorbent paper in an autoclave-safe container and autoclaved for 30 min at 121°C, 100 kiloPascals. Autoclaved coupons were stored in sterile containers until used. Microbial recovery from autoclaved APC coupons was variable. There was no visible damage, changes in contact angle measurements, or evidence of surface variability on autoclaved APC coupons after inspection with a scanning electron microscope (data not shown). In order to clean and remove interferents such as dried, residual paint solvent, APC coupons were soaked in 70% ethanol for 20 min, rinsed at least 3 times with sterile 18 mega-ohm-cm water and dried.

### Response surface methodology (RSM)

RSM was used for previous work with HHA decontamination of materials contaminated with spores because the RSM Design of Experiments (DOE) was requested by the funding agencies as an industry-accepted method to describe HHA points of failure (Beauregard 1992; Lenth 2009; Montgomery 2009; Myers et al. 2009; Buhr et al. 2012, 2015). RSM required three equally spaced test parameters for each variable; time, temperature, and humidity. An extensive amount of trial-and-error testing showed a very rapid and dramatic impact of temperature and RH on virus decontamination. Test temperatures were 55, 60, and 65°C. Test RH was 50, 70, and 90%. Test times were 1, 5, and 9 h to provide a practical timeframe for decontamination of ≤24 h. RSM DOE test parameters are displayed in **Figure 1**. The first 13 test runs (first iteration) included the center and corners of the experimental test box. The latter 6 test runs (second iteration) included the face-points at the center of each side of the test box.

**Figure 1.**
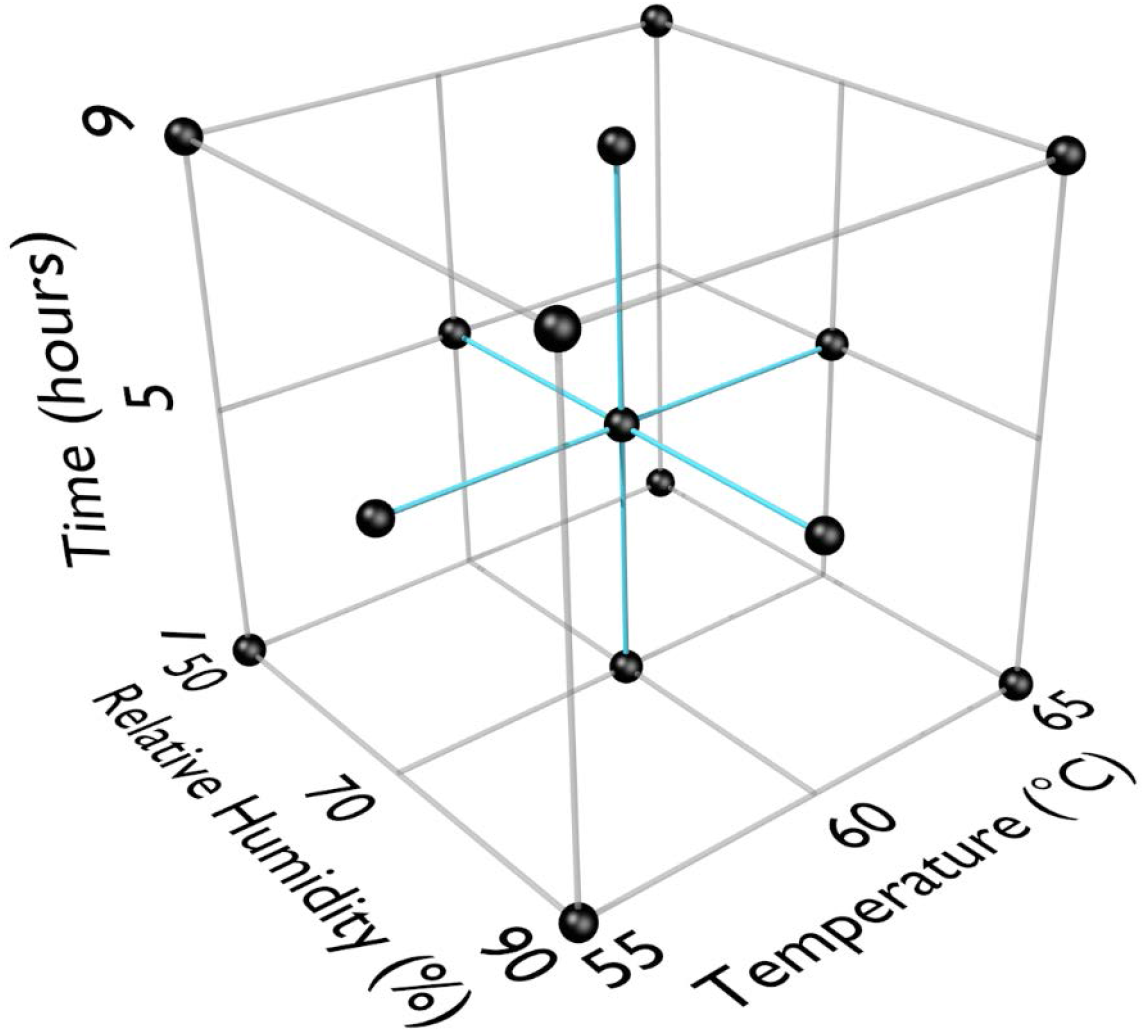
Response Surface Methodology (RSM) Design of Experiments (DOE) for three test factors (°C, % relative humidity, time in h).

### Virus Test Method and Design

An overlay procedure used for Φ6 was adapted from procedures used for *Escherichia coli* and its bacteriophages (Sambrook et al. 1989) using TSA and TSB media instead of Luria-Bertani agar and broth. Plates were incubated at 26±2°C for 20±2 h before quantifying plaques. Plates were then incubated another 24±2 h at RT and scored a final time.

Φ6 stock preps were removed from −80°C storage and thawed at RT prior to preparing inoculum by transferring stock Φ6 into 50-ml conical tubes containing 10mM HEPES + 10% Sucrose pH 7.0 for a final concentration of 1-2e9 PFU ml^-1^. Test coupons and conical tubes were inoculated with 0.1 ml of Φ6 inoculum, held at RT for 24±2 h to dry allowing virus to adhere to the test material, then transferred to TPP^®^ tubes. Following exposure of Φ6 to various hot humid air decontamination testing parameters, samples were extracted and plated.

For Φ6 extraction, 5 ml of 10mM HEPES + 10% sucrose pH7 were added to each conical tube with a virus-inoculated coupon and vortexed for 2 min. After vortexing, 5 ml of HB10Y log-phase culture were added and allowed to infect at RT for 15 min, followed by 2 min of vortexing. Each sample was serially diluted in 900 ul of 10 mM HEPES + 10% Sucrose pH7 out to the −6 dilution. For each Φ6 dilution, 1,000 ul for the first dilution and 200 ul for subsequent dilutions were transferred into individual tubes containing 200 ul log-phase HB10Y. Then 1,200 ul or 200 ul of those Φ6/HB10Y mixtures were added to individual TSB overlay tubes, poured onto individual TSA plates and allowed to solidify for ≥30 min. Solidified plates were then inverted, incubated for 20+/-2 h at 26°C and quantified. Plates were incubated another 24 h at RT and quantified a final time. Quantitation and calculations of survival were performed as previously described (Buhr et al. 2012, 2014). The maximum (100%) recoverable virus reference, the inoculum titer, was used to calculate extraction efficiency. Extraction efficiency was the number of PFU removed from non-treated control coupons divided by the number of PFU in the inoculum.

### Response Surface Methodology (RSM) Performance Envelopes - Mathematical Model Development

RSM to represent survivability of Φ6 bacteriophage were developed. For a given Design of Experiment (DOE) point, sample data varied with respect to initial Φ6 bacteriophage population and a final surviving Φ6 bacteriophage population. To facilitate model development, the results of the samples were normalized into a Survival Fraction (SF). A Survival Fraction is simply the surviving population divided by the initial population. Survival Fraction is assumed to never be greater than one or less than zero.

The smallest non-zero Survival Fraction (SF) that may be observed is limited due to the initial population being a finite number. Specifically, the smallest observable SF is 1 divided by the initial population. However, if the results reflect that of a 1 ml sample taken from a 10 ml sample (as is true for this data), then the smallest observable SF is 10 divided by the initial population. If extraction efficiency is less than 100%, then the smallest observable SF becomes 10 divided by the initial population divided by the extraction efficiency. If test data shows a surviving population of zero, it is likely because the theoretical SF is smaller than the smallest observable SF. Because of the limitation imposed by a finite initial population, any test result showing a survival population of zero was treated as a SF that is less than or equal to the smallest observable SF.

The geometric mean survival fraction (GMSF) equation below shows how GMSF was calculated for each DOE point. In this equation, “N” represents the number of samples for a given DOE point.

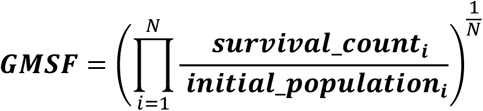

All GMSF values from the experiments lie between “0” and “1”, and they differ from each other by orders of magnitude, which can make it difficult to find a good fit to the data. To avoid this problem and to facilitate the data fitting process, it is desirable to transform the widely varying GMSF values such that the transformed values are distributed in a linear fashion. To achieve this, the natural log of the GMSF values was computed. However, this transform still resulted in a very non-linear distribution. A second transform was performed, using Equation 1 below, that flipped the sign of the first transform and then took its natural log. This yielded a more linear distribution of data.

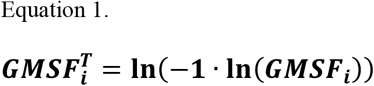

One benefit of this transform is that it constrains all model predictions to a value between “0” and “1” and thereby prevents a result that contradicts the definition of SF. Data fit exploration focused on obtaining a good fit to the data set of transformed GMSF values. During the data fitting process, a constraint was imposed on the fit such that as temperature and/or RH and/or time increase, SF decreases consistently without exception. The final functional form for predicting GMSF is shown in the system of equations below (Equation 2.)

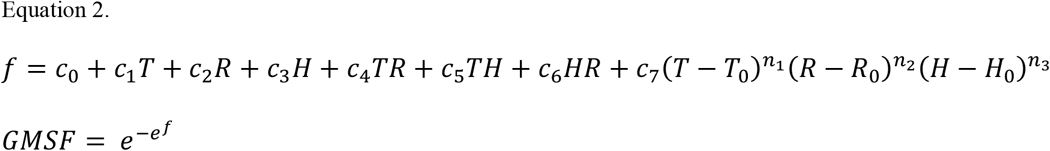

Where:

*T = Temperature (°C), R = Relative Humdity (%), H = Time (hours)*
*c_0_ to c_7_ = addition and multiplication coefficients*
*n_1_ to n_3_ = power coefficients*
*T_0_ = Temperature offset coefficent*
*R_0_ = Relative Humdity offset coefficient*
*H_0_ = Time offset coefficient*
*GMSF = Geometric Mean Survival Fraction*

As is evident in the data, a given exposure condition will result in a distribution of various SF values that can vary geometrically. Because of this geometric variation, it is appropriate to assume the natural log of the survival fractions for a given sample follow a Normal distribution for a given exposure condition. An estimate of the expected distribution of results is important because it helps a user identify a probability that an expected total kill will occur for a given exposure condition. For example, a user can utilize the expected distribution to identify the exposure condition required to achieve at least a 7-log_10_ kill with a 95% probability.

Most of the exposure DOE points consist of only 5 samples and this small sample size increases the uncertainty in determining the population Geometric Standard Deviation (GSD) from the sample geometric standard deviation. There was no clear and consistent pattern in the sample geometric standard deviations that would allow for the development of a reasonable equation to predict geometric standard deviation as a function of exposure conditions. However, the DOE center point (Temp = 60, RH = 70, H = 5) consists of 25 samples for each medium and consists of enough data to reasonably estimate a geometric standard deviation that is assumed to be representative for the entire DOE factor space. Using the test results from the DOE center point for each medium, the geometric standard deviation was computed using Equation 3 below.

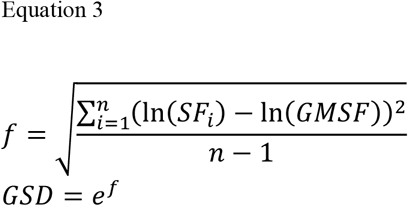

Where:

*n = Number of samples*
*i = indexed reference to a particular sample* *SF_i_ = Survival Fraction for sample “i”*
*GMSF = Geometric Mean Survival Fraction*
*GSD = Geometric Standard Deviation*

The Normal distribution of possible Survival Fraction logs for a given exposure condition is described by the natural log of the GMSF predicted using Equation 2 and the natural log of the GSD computed using Equation 3. The GMSF will vary based on exposure conditions and medium but the GSD will only vary by medium. The equations for the Normal distribution are provided in Equations 4 and 5 below.

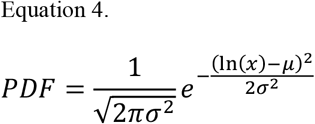

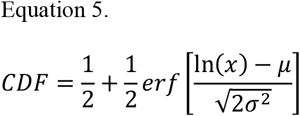

Where:

*PDF=Probability Distribution Function. Characterizes the relative frequency of occurrence*.
*CDF=Cumulative Distribution Function. The integral of the PDF with results ranging from 0 to 1*.
*x = Horizontal axis of possible Survival Fractions*
*μ = Natural log of Geometric Mean Survival Fraction (GMSF)*
*σ = Natural log of Geometric Standard Deviation (GSD)*
*erf = The error function*

The Cumulative Distribution Function (CDF) equation returns the area for a region under the Probability Distribution Function (PDF) curve by integrating from −∞ to a specified SF. A CDF result of “0.95” indicates that 95% of the SF values observed will be less than the specified SF.

The Solver add-in for Microsoft Excel 2016 was used to explore various fits to the transformed survival fractions. The Solver add-in has the added benefit of allowing fit constraints which was crucial to obtaining a good and reasonable fit. The “Evolutionary” method in Solver proved a valuable tool in optimizing the coefficients shown in Equation 2. Excel was also used to produce the various plots in this report and assess model performance.

### C-130 and C-17 Aircraft Test Demonstrations

Enveloped virus test indicators consisted of APC coupons inoculated with 1 of 5 independent virus preparations at >8 log_10_ coupon^-1^ and the test coupons were placed inside TPP^®^ tubes prior to storage, shipping and testing. Test indicators were placed in 20-22 locations with 5 test indicators per location within the aircraft. Each of the 5 test indicators was inoculated with an independent virus preparation. HHA tests were conducted inside a C130 fuselage located at Army Combat Capabilities Developmental Command, Edgewood, MD, USA (CCDC) (**Figure 2**) or in a C-17 located at Dover Air Force Base, Dover, DE, USA (**Figure 3**) using a previously tested HHA system (Buhr et al, 2016) ran and managed/monitored by a consortium listed in the acknowledgements. After HHA exposure, the test indicators were shipped to a laboratory for virus extraction. Shipping controls were shipped, but not treated with HHA. Laboratory controls were not shipped. Both control sets were prepared, extracted and quantified alongside the test indicators.

**Figure 2.**
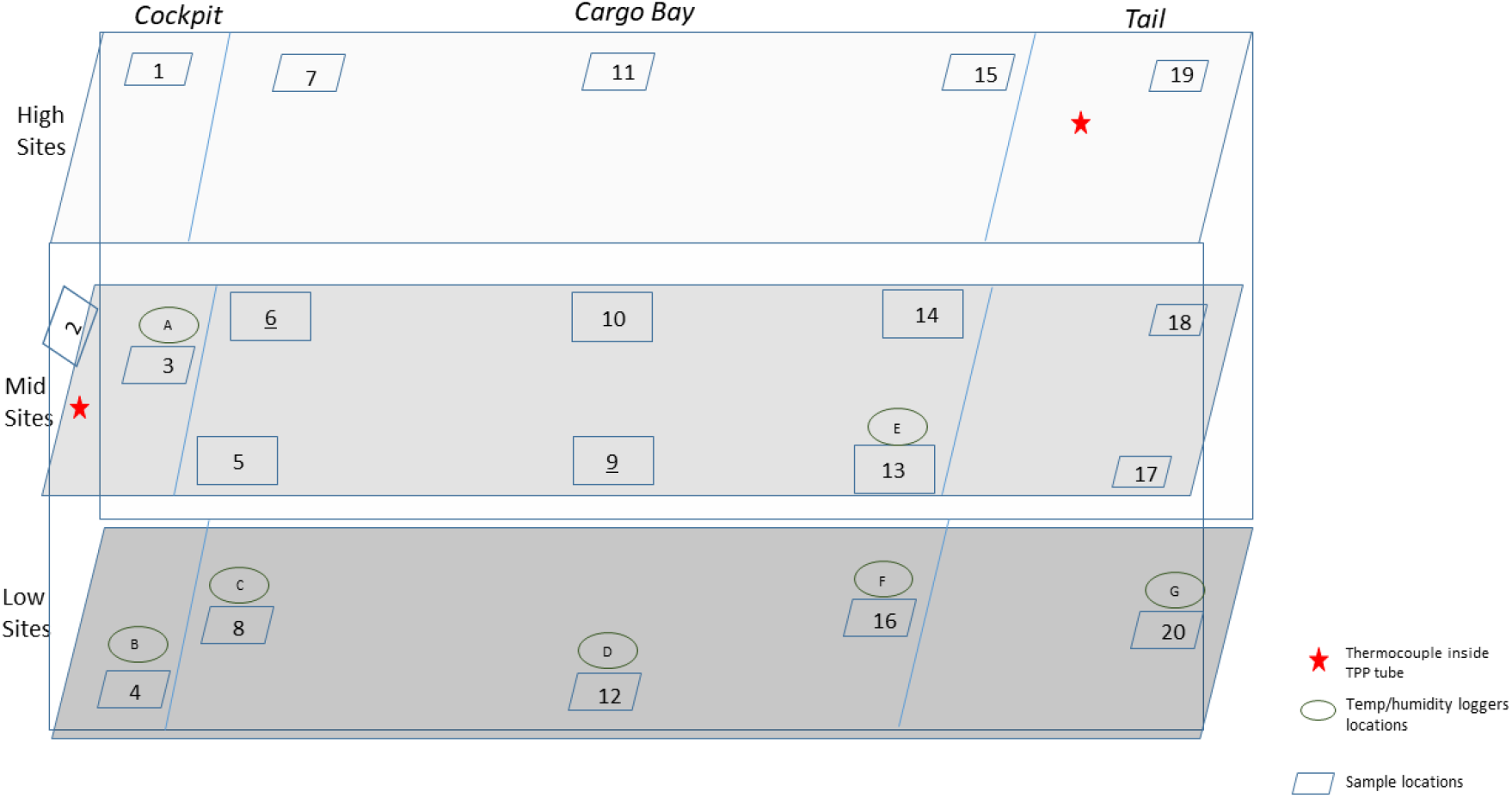
Side view of a C-130 showing the sampling layout for low, middle and high sampling sites.

## RESULTS

### HHA Inactivation of Enveloped RNA Virus

In order to develop RSM performance envelopes, partial inactivation data was needed especially at the DOE mid-point. A small test at 50, 60, 70 and 80°C, RH at 5% or 90% for 1 and 24 h produced critical data showing that high RH was critical for enveloped virus inactivation (**Table 1**). There was only 2.4 log_10_ enveloped virus inactivation at 70°C, 5% RH for 24 h compared to complete inactivation at 70°C, 90% RH for 1 h. In addition, there was no enveloped virus inactivation after drying enveloped virus on NTC and treating at 26.7°C at 5, 30, 55 and 80% RH for up to 10 d (data not shown). These data drove the test parameters shown in **Figure 1**. Although 65°C was greater than the original 60°C temperature goal, 65°C was selected for the experimental design because inactivation at <60°C was too slow to generate RSM that met the <24 h objective. Therefore, 65°C was incorporated in the DOE and 60°C, 70% RH, 5 h was set as the mid-point for the final DOE in order to obtain sufficient partial kill data to develop the RSM performance envelopes.

**Table 1.** Log_10_ PFU recovered (survival) and reduction (L.R.) after drying Φ6 on APC coupons (enveloped virus test indicator) and then hot air exposure at 5% or 90% RH for 1 or 24 h. Five test indicators per test. Not Applicable (NA).

| Parameters | Survival | L.R. | Parameters | Survival | L.R. |
| --- | --- | --- | --- | --- | --- |
| 50°C, 5%, 1 h | 7.5 ± 0.2 | 0.8 ± 0.1 | 50°C, 90%, 1 h | 7.3 ± 0.4 | 0.9 ± 0.2 |
| 60°C, 5%, 1 h | 7.0 ± 0.3 | 1.3 ± 0.2 | 60°C, 90%, 1 h | 1.4 ± 0.3 | 6.8 ± 0.1 |
| 70°C, 5%, 1 h | 6.3 ± 0.2 | 2.0 ± 0.1 | 70°C, 90%, 1 h | 0.0 ± 0.0 | 8.2 ± 0.0 |
| 80°C, 5%, 1 h | 6.1 ± 0.2 | 2.2 ± 0.1 | 80°C, 90%, 1 h | 0.0 ± 0.0 | 8.2 ± 0.0 |
| Inoculum - 1 h | 8.3 ± 0.0 | NA | Inoculum - 1 h | 8.2 ± 0.1 | NA |
| 50°C, 5%, 24 h | 6.8 ± 0.4 | 1.4 ± 0.2 | 50°C, 90%, 24 h | 1.1 ± 1.6 | 6.9 ± 0.7 |
| 60°C, 5%, 24 h | 6.2 ± 0.2 | 2.0 ± 0.1 | 60°C, 90%, 24 h | 0.0 ± 0.0 | 8.0 ± 0.0 |
| 70°C, 5%, 24 h | 5.8 ± 0.5 | 2.4 ± 0.2 | 70°C, 90%, 24 h | 0.0 ± 0.0 | 8.0 ± 0.0 |
| 80°C, 5%, 24 h | 1.4 ± 1.3 | 6.8 ± 0.6 | 80°C, 90%, 24 h | 0.0 ± 0.0 | 8.0 ± 0.0 |
| Inoculum - 24 h | 8.2 ± 0.1 | NA | Inoculum - 24 h | 8.0 ± 0.1 | NA |

Overall, there were several thousand trial-and-error data points leading to the HHA decontamination DOE data set in **Table 2**, which shows data from nineteen test runs with five independent virus preparations per coupon type. There were 475 Φ6 virus test samples and 475 corresponding controls, plus non-inoculated coupon controls. Each test variable (temperature, RH and time) had an impact on decontamination. Importantly, high RH, high temperature and longer times increased inactivation rates as expected. Inactivation of enveloped virus dried on to materials was measured at ≥7 log_10_ at 60°C, 90%, for 9 h. This surpassed the original goal of ≥7 log_10_ at ≤6O°C, 24 h. Furthermore, enveloped virus inoculum held in aqueous solution (solution controls) were shown to be much more sensitive to the rise in temperature than virus dried on to materials and subjected to hot, humid air. Therefore, water and/or high humidity were significant parameters for inactivating enveloped virus.

**Table 2.** Log_10_ virus survival of Φ6 after drying on to different materials and then exposure to different temperatures, RH and time according to the DOE in Figure 1. The DOE consisted of 13 independent runs - 8.4±0.0 log_10_; 6 independent runs - 8.3±0.1 log_10_; 5 independent tests per material per test run. APC – Aircraft Performance Coating; Solution Control – wet virus inoculum – not dried on a material; NA – not tested for DOE. Hours (h). Polypropylene material from the TPP^®^ tubes was the material. *An outlier was removed.

|  |  | 50% RH | 50% RH | 50% RH | 70% RH | 70% RH | 70% RH | 90% RH | 90% RH | 90% RH |
| --- | --- | --- | --- | --- | --- | --- | --- | --- | --- | --- |
| **Material** | °C | 1 h | 5 h | 9 h | 1 h | 5 h | 9 h | 1 h | 5 h | 9 h |
| **Nylon** | 55 | 8.2 ± 0.2 | NA | 7.0 ± 0.2 | NA | 5.5 ± 0.1 | NA | 7.2 ± 0.1 | NA | 1.1 ± 1.5 |
| **Nylon** | 60 | NA | 7.0 ± 0.4 | NA | 6.7 ± 0.2 | 4.9 ± 0.6 | 3.4 ± 0.3 | NA | 0.0 ± 0.0 | NA |
| **Nylon** | 65 | 7.3 ± 0.2 | NA | 5.6 ± 0.1 | NA | 0.7 ± 0.9 | NA | 0.0 ± 0.0 | NA | 0.0 ± 0.0* |
| **Polypropylene** | 55 | 8.3 ± 0.3 | NA | 7.6 ± 0.2 | NA | 7.8 ± 0.3 | NA | 7.7 ± 0.2 | NA | 0.0 ± 0.0 |
| **Polypropylene** | 60 | NA | 6.9 ± 0.2 | NA | 8.1 ± 0.1 | 6.5 ± 0.7 | 4.9 ± 0.3 | NA | 0.0 ± 0.0 | NA |
| **Polypropylene** | 65 | 7.2 ± 0.2 | NA | 2.4 ± 0.4 | NA | 2.0 ± 0.3 | NA | 0.0 ± 0.0 | NA | 0.0 ± 0.0 |
| **Wiring** | 55 | 8.3 ± 0.1 | NA | 7.9 ± 0.3 | NA | 7.0 ± 0.8 | NA | 7.5 ± 0.1 | NA | 0.0 ± 0.0 |
| **Wiring** | 60 | NA | 6.9 ± 0.6 | NA | 7.9 ± 0.1 | 6.7 ± 0.5 | 4.3 ± 0.5 | NA | 0.0 ± 0.0 | NA |
| **Wiring** | 65 | 7.0 ± 0.4 | NA | 4.1 ± 0.6 | NA | 2.0 ± 1.4 | NA | 0.0 ± 0.0 | NA | 0.0 ± 0.0 |
| **APC** | 55 | 8.3 ± 0.1 | NA | 6.8 ± 0.4 | NA | 7.0 ± 0.6 | NA | 7.5 ± 0.2 | NA | 1.3 ± 1.2 |
| **APC** | 60 | NA | 3.8 ± 0.9 | NA | 7.8 ± 0.2 | 6.2 ± 0.7 | 4.6 ± 0.3 | NA | 0.0 ± 0.0 | NA |
| **APC** | 65 | 5.3 ± 0.7 | NA | 2.9 ± 1.6 | NA | 0.8 ± 0.8 | NA | 0.0 ± 0.0 | NA | 0.0 ± 0.0 |
| **Solution Control** | 55 | 0.0 ± 0.0 | NA | 0.3 ± 0.6 | NA | 0.0 ± 0.0 | NA | 0.4 ± 0.9 | NA | 0.0 ± 0.0 |
| **Solution Control** | 60 | NA | 0.0 ± 0.0 | NA | 0.0 ± 0.0 | 0.0 ± 0.0 | 0.0 ± 0.0 | NA | 0.0 ± 0.0 | NA |
| **Solution Control** | 65 | 0.0 ± 0.0 | NA | 0.0 ± 0.0 | NA | 0.0 ± 0.0 | NA | 0.0 ± 0.0 | NA | 0.0 ± 0.0 |

### HHA RSM for Enveloped RNA Virus Inactivation

RSM for each material (**Figure 4**) were constructed from the DOE Φ6 inactivation data in **Table 2**. Inactivation of Φ6 virus was similar on all coupon types indicating that material type had little impact on inactivation kinetics. Since virus inactivation was similar across each material an additional composite RSM was generated (**Figure 4**). The models more clearly showed virus inactivation kinetics compared to the table data. At 1 h, there was a 90% probability of at least a 7-log_10_ virus inactivation at 65°C, 90% RH after virus-contaminated materials were treated with HHA.

The composite model after 1 h of virus inactivation was critical for directing HHA parameters during the aircraft field-testing. The composite models were used to translate HHA parameters to dew points/absolute humidity levels (data not shown). The translation made it apparent that absolute humidity or dew point strongly correlated with inactivation. For example, there was a 90% probability of a 4-log_10_ kill at ≥61.7°C and a dew point humidity ≥59.4°C. The temperature could go up or down with the same inactivation kinetics so long as the absolute humidity/dew point was maintained. Hence, the inactivation kinetics correlated with the total amount of water in the air. Models with dew points and/or absolute humidity are in the queue for future development.

### C-130 Aircraft Field Test

A total of 100 enveloped virus test indicators were distributed at 20 sites throughout a C-130 fuselage as depicted in **Figure 2**. Based on the RSM in **Figure 4** and with a >4 log_10_ inactivation goal, the HHA goals for the C-130 were ≥61.7°C and a dew point humidity of ≥59.4°C for 1 h, excluding ramp-up and ramp-down times. The measured HHA parameters for the C-130 are shown in **Figure 5**. As shown, the temperature and humidity goals were exceeded. In addition, the ramp-up time was 2.5-3 h, which was longer than lab testing with a ramp-up of 1 h, hence ramp-up conditions on the aircraft likely contributed to enveloped virus inactivation. Ramp-down time was nearly 2 h, which was also much longer than lab conditions. The total HHA time including ramp-up and ramp-down for the C-130 was 5.5 h compared to 2 h in the lab-based RSM. The C-130 temperatures were always above the dew points so there was no condensation on the virus-inoculated APC samples (**Figure 5**). Samples were shipped to a lab for extraction. **Table 3** shows 8.1±0.1 log_10_ PFU were recovered on 10 non-shipped control samples and 25 ground-shipped control samples out of 8.3±0.1 log_10_ PFU dried on to the APC coupons. The loss of 0.2 log_10_ PFU on the control coupons could have been due to extraction efficiency and/or a small drop in virus viability. In either case, the 8.1 log_10_ PFU recovery on the control coupons was high and it validated there was no significant loss of viability during transportation to and from the field site. The limit of detection was 1.0 log_10_, and since there was no recovery from test indicators the log reduction can confidently be reported as 7.1 log_10_ (8.1 – 1.0 log_10_). Since the temperature and RH limits were exceeded for a 7-log_10_ kill and there was no viable virus recovered from any test indicator, there is practical confidence that a complete >8-log_10_ kill was achieved.

**Table 3.** Log_10_ PFU recovered (survival) and reduction (L.R.) after drying 8.3±0.1 log_10_ of Φ6 per APC coupon (enveloped virus test indicator) and then HHA exposure in a C-130 aircraft. The limit of detection was 1.0 log_10_ and the detection range was 1.0-8.1 log_10_ where the upper limit was the amount extracted from the controls. Since there was no enveloped virus recovery on any test indicator there was practical confidence of a complete virus kill. The number of test indicators are in parentheses. Not Applicable (NA).

| Test Indicators | Survival | L.R. |
| --- | --- | --- |
| Non-Shipped Controls (10) | 8.1±0.1 | NA |
| Shipped Controls (25) | 8.1±0.1 | NA |
| Test Indicators (100) | 0 | 8.1±0.1 |

### C-17 Aircraft Field Test

A total of 110 enveloped virus test indicators were distributed throughout the C-17 as depicted in **Figure 3**. Based on the RSM in **Figure 4**, results from the C-130 test, and with a >4 log_10_ inactivation goal, the HHA parameters were set to ≥61.7°C and a dew point humidity of ≥57.2°C for the C-17, at which point a 1 h “hold time” was started. Another goal was to increase the temperature and humidity during the 1 h “hold time” to 61.7°C and a dew point humidity of ≥59.4°C. The measured HHA parameters for the C-17 are shown in **Figure 6**. The temperature reached 65.0°C and humidity reached a 60.6°C dew point, surpassing the temperature and humidity goals during the 1-h hold time. In addition, the ramp-up time was 1.5 h, which saved >1 h compared to the C-130 test, but was 0.5 h longer than lab tests. Ramp-down time was 45 min compared to 2 h for the C-130 and only one min for lab tests. The total HHA time for the C-17 was <3.5 h compared to 5.5 h for the C-130 and 2 h for lab tests. The C-17 temperatures were always above the dew points so there was no condensation on the virus-inoculated APC samples (**Figure 6**). Samples were ground-shipped to a lab for extraction. **Table 4** shows 7.6±0.1 log_10_ PFU on 10 non-shipped control samples and 25 ground-shipped control samples out of 8.2±0.1 log_10_ PFU dried on to the APC coupons. The loss of 0.6 log_10_ PFU on the control coupons could have been due to extraction efficiency and/or a drop in virus viability. In either case, the 7.6 log_10_ PFU recovery on the control coupons was high and it showed no practical significant loss of virus viability during transportation to and from the field site. The limit of detection was 1.0 log_10_. Since there was viable PFU from at least one of the 110 test indicators, all samples with no viable PFU were assigned 1.0 log_10_. Site 15 on the C-17 showed the lowest log reduction of 5.9±0.6 log_10_ PFU. This site had the most air movement. Although there were no TPP^®^ tubes with temperature and RH readings at Site 15, we speculate that the rapid air movement slowed the penetration of moist air across the 0.2 um membrane of the TPP^®^ tubes, hence the lowest level of inactivation. Every other C-17 location had at least a 6-log_10_ reduction. Ramp-up/ramp-down times were longer than the lab tests used for RSM, and raising the heat and humidity during the 1 h hold time contributed to the virus inactivation. Hence the goal of >4 log_10_ reduction was surpassed as predicted by RSM.

**Figure 3.**
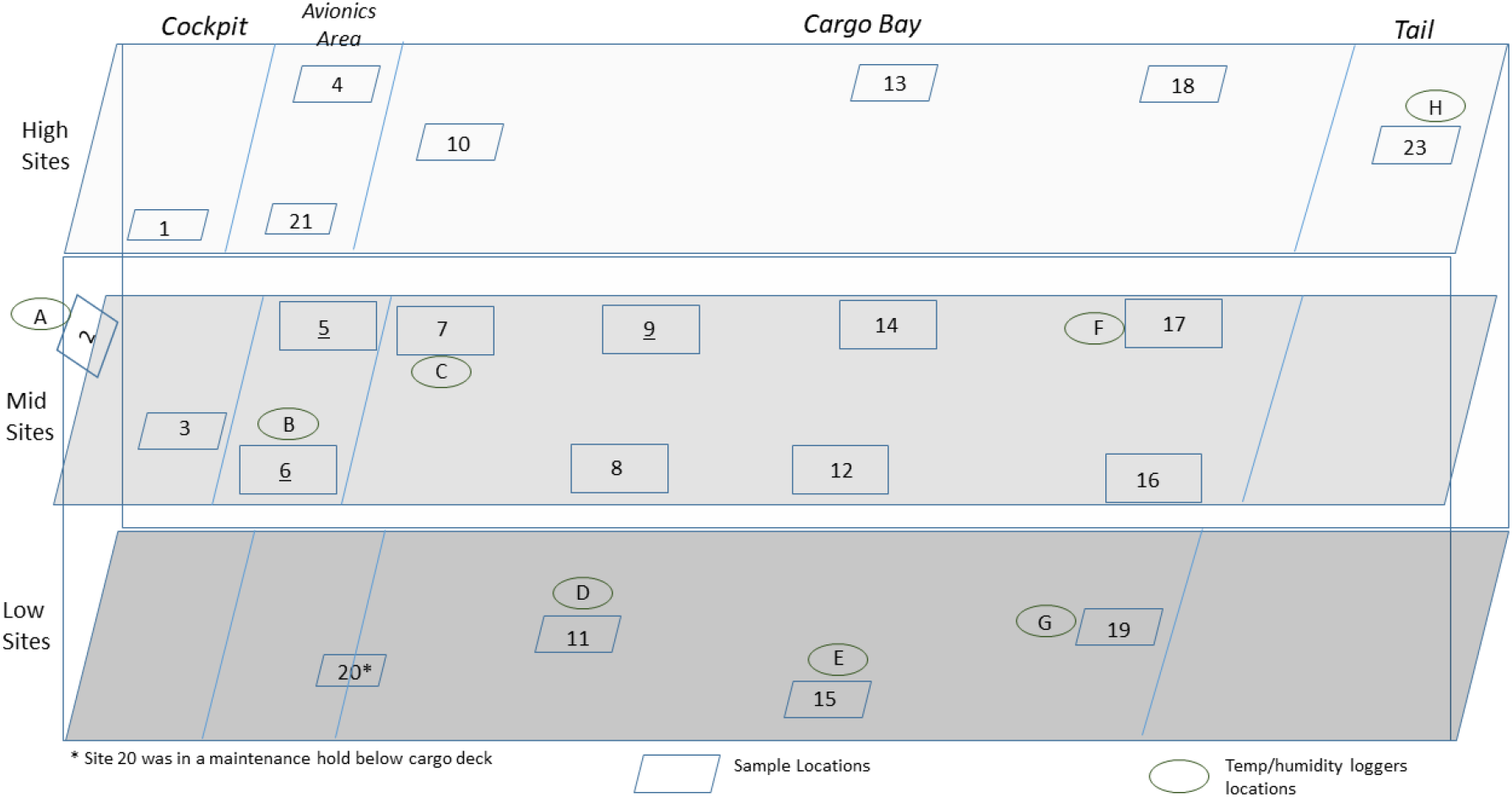
Side view of a C-17 showing the sampling layout for low, middle and high sampling sites.

**Figure 4.**
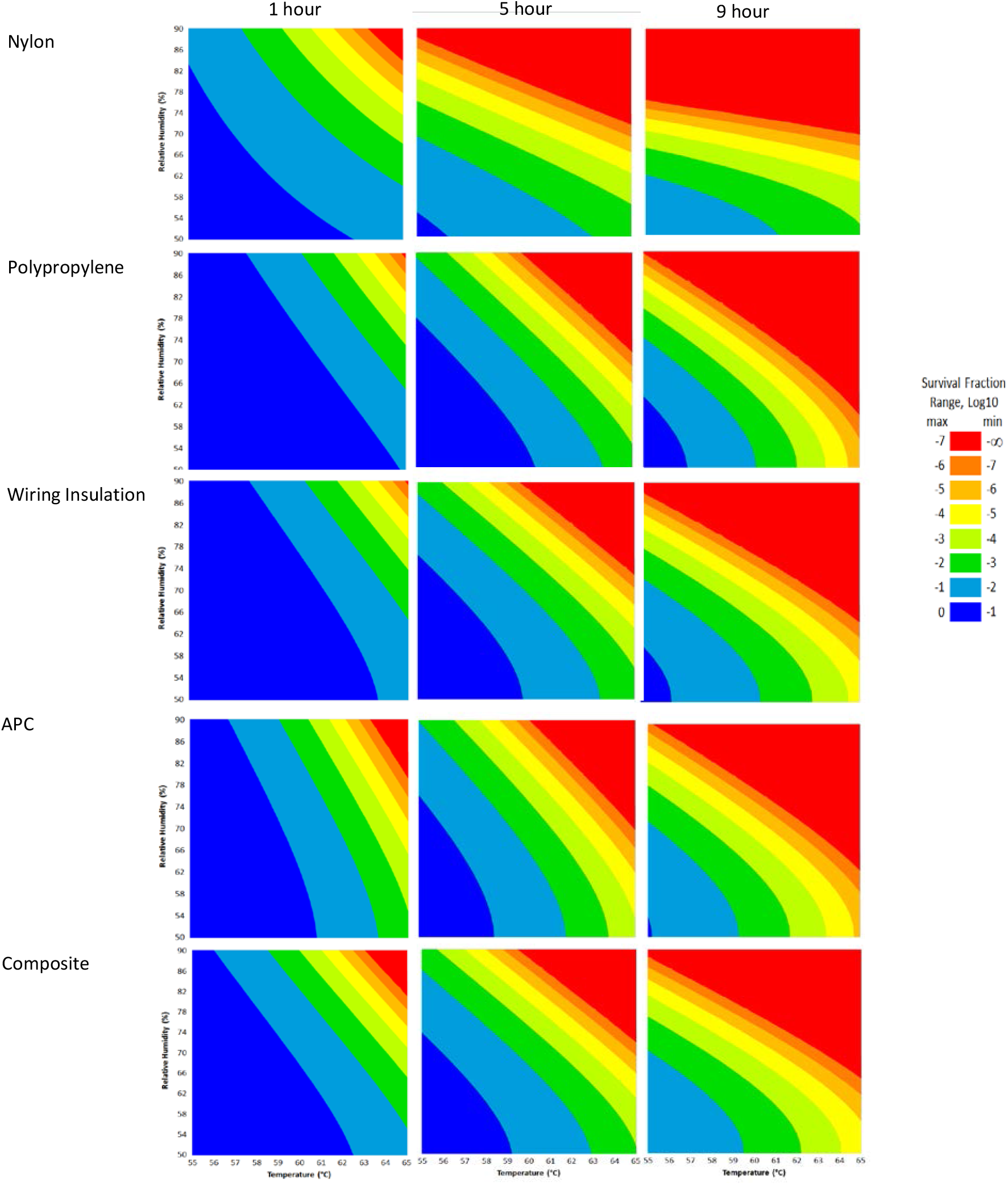
Models with a 90% statistical probability of Φ6 virus inactivation after virus was dried on to nylon, polypropylene, wiring insulation, or aircraft performance coating after incubation in hot humid air. Composite models are the accumulation of data for all 4 materials. Areas are represented as >7-log_10_ (red), 6-7 log_10_ (orange), 5–6 log_10_ (gold), 4–5 log_10_ (yellow), 3–4 log_10_ (green), 2–3 log_10_ (dark green), 1–2 log_10_ (blue), <1 log_10_ (dark blue) virus inactivation.

**Figure 5.**
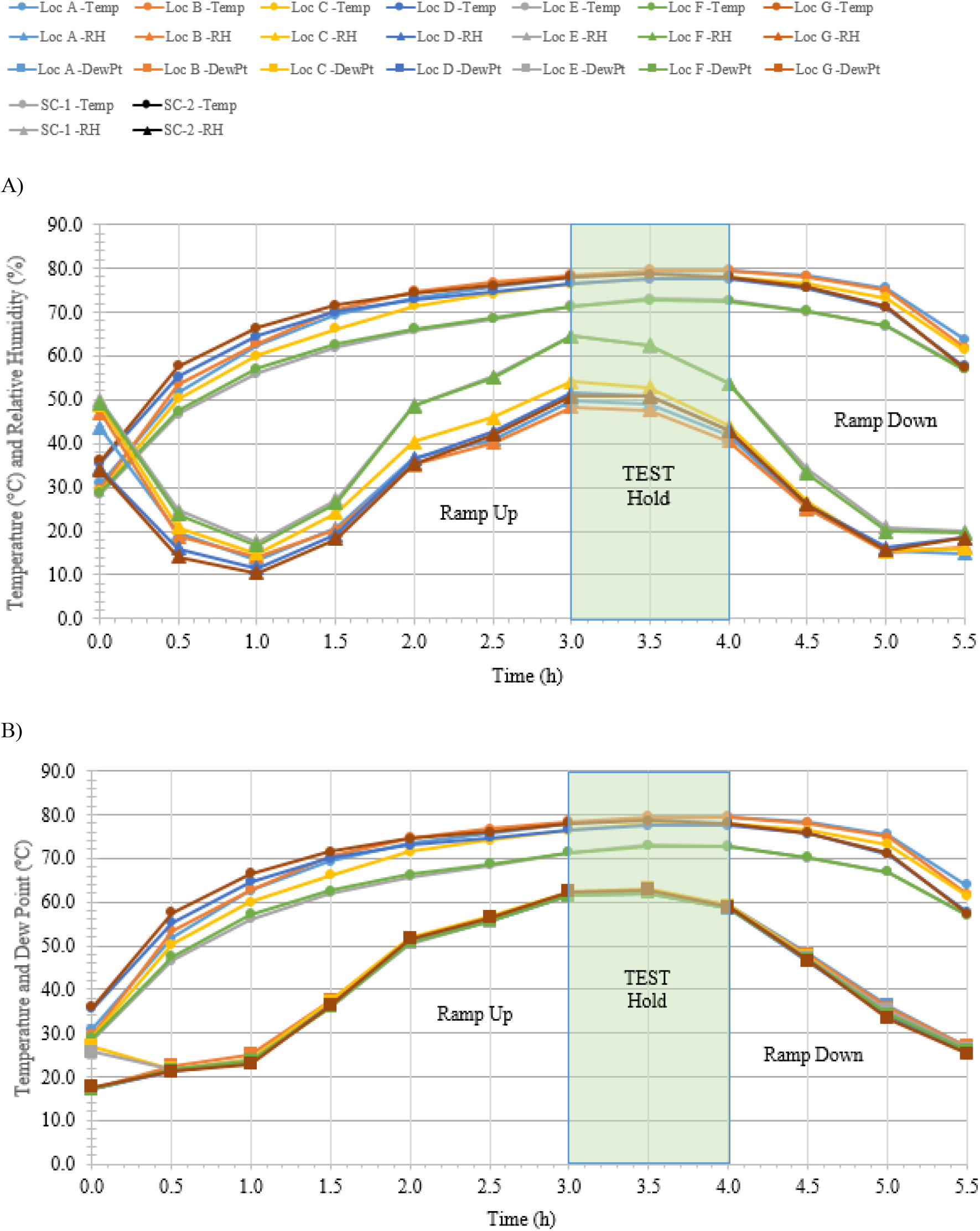
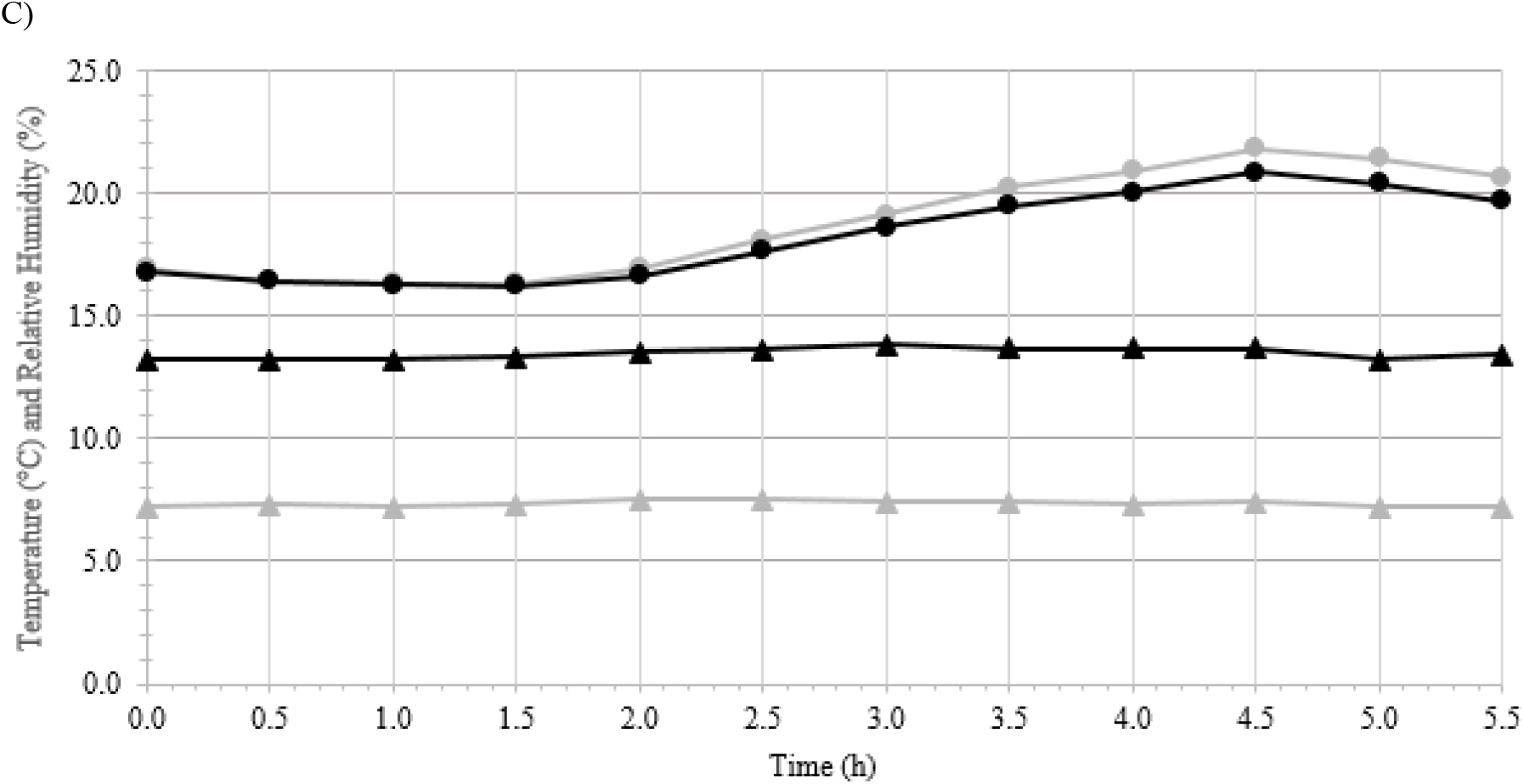
Temperature and humidity profiles during HHA decontamination of a C-130. A) Temperature (°C) and RH (%) from seven locations (A-G) throughout the C-130. B) Temperature and dew point (°C) from seven locations (A-G) throughout the C-130. C) Temperature (°C) and RH (%) logged within two sets of shipping controls (SC).

**Figure 6.**
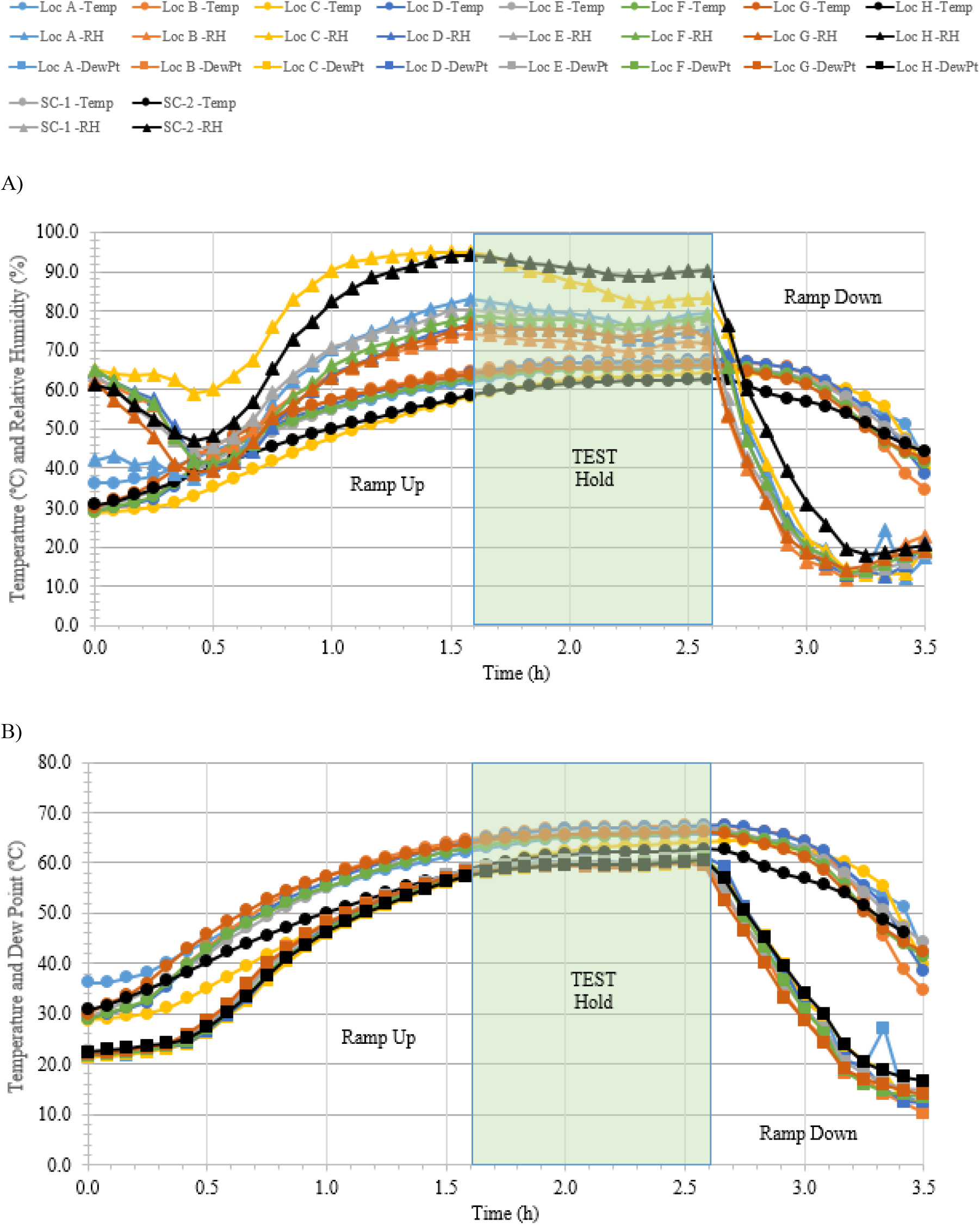
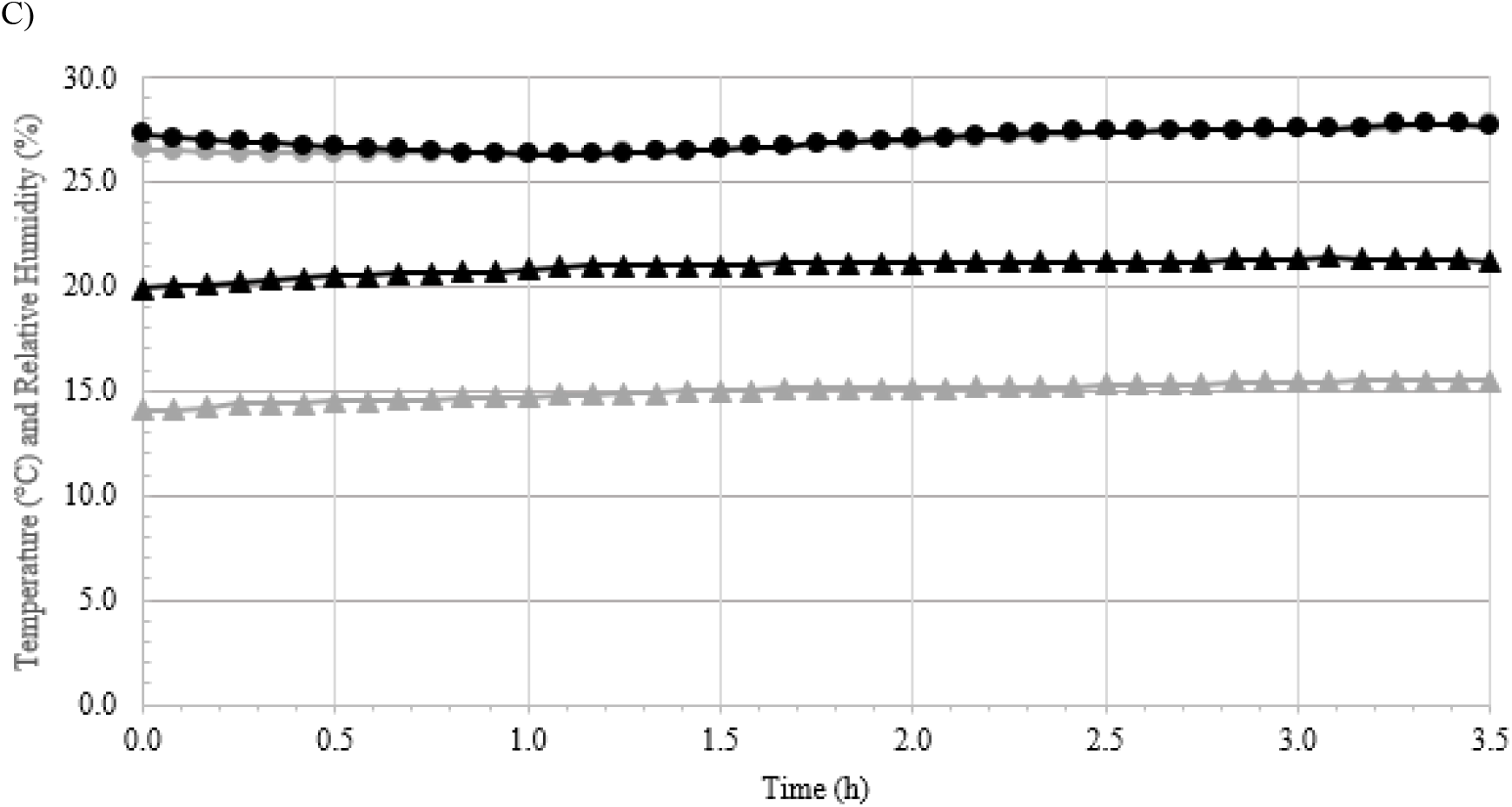
Temperature and humidity profiles during HHA decontamination of a C-17. A) Temperature (°C) and RH (%) from eight locations (A-H) throughout the C-17. B) Temperature and dew point (°C) from eight locations (A-H) throughout the C-17. C) Temperature (°C) and RH (%) logged within two sets of shipping controls (SC).

**Table 4.** Log_10_ PFU recovered (survival) and reduction (L.R.) after drying 8.2±0.1 log_10_ of Φ6 per APC coupon (enveloped virus test indicator) and then HHA exposure in a C-17 aircraft. The limit of detection was 1.0 log_10_ and the detection range was 1.0-7.6 log_10_ where the upper limit was the amount extracted from the controls. Since there was enveloped virus recovery on some test indicators the lowest recorded kill level was 1.0 log_10_ even if there were no viable PFU measured from a test indicator. The number of test indicators are in parentheses. Not Applicable (NA).

| Test Indicator Locations | Survival | L.R. |
| --- | --- | --- |
| Non-Shipped Controls (10) | 7.6 ± 0.1 | NA |
| Shipped Controls (25) | 7.6 ± 0.1 | NA |
| 22 - Repurposed As a Control On-Site (5) | 7.6 ± 0.2 | NA |
| 1 - Flight Deck (FD) - Ceiling (5) | 1.4 ± 0.8 | 6.2 ± 0.3 |
| 2 - FD - Glare shield (5) | 1.3 ± 0.6 | 6.3 ± 0.3 |
| 3 - FD - Crew rest area (5) | 1.0 ± 0.0 | 6.6 ± 0.1 |
| 4 - Avionics - Rack doghouse (5) | 1.0 ± 0.0 | 6.6 ± 0.1 |
| 5 - Lower avionics rack (5) | 1.0 ± 0.0 | 6.6 ± 0.1 |
| 6 - Lavatory (5) | 1.5 ± 0.9 | 6.1 ± 0.4 |
| 7 - Load master station (5) | 1.3 ± 0.5 | 6.3 ± 0.2 |
| 8 - Forward Cargo Bay (FCB) - Left (5) | 1.5 ± 0.9 | 6.1 ± 0.4 |
| 9 - FCB - Right (5) | 1.4 ± 0.7 | 6.2 ± 0.3 |
| 10 - FCB - Ceiling (5) | 1.3 ± 0.6 | 6.3 ± 0.3 |
| 11 - FCB - Floor (5) | 1.4 ± 0.9 | 6.2 ± 0.4 |
| 12 - Mid Cargo Bay (MCB) - Left (5) | 1.5 ± 0.9 | 6.1 ± 0.4 |
| 13 - MCB - Right (5) | 1.0 ± 0.0 | 6.6 ± 0.1 |
| 14 - MCB - Ceiling (5) | 1.3 ± 0.7 | 6.3 ± 0.3 |
| 15 - MCB - Floor (5) | 1.7 ± 1.3 | 5.9 ± 0.6 |
| 16 - Aft Cargo Bay (ACB) - Left (5) | 1.0 ± 0.0 | 6.6 ± 0.1 |
| 17 - ACB - Right (5) | 1.4 ± 0.8 | 6.2 ± 0.4 |
| 18 - ACB - Ceiling (5) | 1.4 ± 0.2 | 6.2 ± 0.1 |
| 19 - ACB - Floor (5) | 1.0 ± 0.0 | 6.6 ± 0.1 |
| 20 - Mx Tunnel - Skin Heat Exchanger (5) | 1.0 ± 0.0 | 6.6 ± 0.1 |
| 21 - Mx Tunnel - Middle (5) | 1.4 ± 0.7 | 6.2 ± 0.3 |
| 23 - Ramp Area (5) | 1.1 ± 0.1 | 6.5 ± 0.1 |

## DISCUSSION

Φ6 was selected as a BSL-1, enveloped RNA virus test indicator for both lab and field tests. There was no Φ6 inactivation after drying on different surfaces after a 10 d exposure to 26.7°C at 80% RH, and there was only 2.4 log_10_ inactivation at 70°C 5% RH for 24 h. The cumulative HHA results from this work suggest that Φ6 may be much more stable to temperature and humidity than reported for other enveloped viruses including SARS-CoV-2 (Sizun et al. 2000, Rabenau et al. 2005, Chan et al. 2011, Anonymous – Department of Homeland Security 2020). The difference in inactivation kinetics among enveloped viruses may reside in test method differences. Indeed, a review of three articles describing dried coronavirus stability showed wide variability in test methods and results. Sizun et al. (2000), Rabenau et al. (2005) and Chan et al. (2011) tested similar coronaviruses including dried SARS-CoV, CoV-229E and CoV-OC-43 at different virus concentrations, virus purities, additives such as fetal calf serum, and fomites, but all the tests were at ambient environmental conditions of 21-25°C. Complete virus inactivation ranged from 3 h up to 13 d across these reports, highlighting the significant impact of test methods on the results. Of interest, the tests with the lowest virus titers had the fastest virus inactivation.

Regardless of whether the Φ6 virus was highly stable and/or the test methods were the driver of stability, the Φ6 results met a commonly accepted criterion where the microbial organism selected for decontamination testing must be a challenge organism equal to or greater than the microbe intended for inactivation such as SARS-CoV-2 (Spaulding 1957; McDonnell and Burke 2011; Anonymous, Environmental Protection Agency 2020). As an example, the United States Environmental Protection Agency (EPA) N-list for decontaminants has been a primary reference point used for decontaminant selection during the COVID19 pandemic (Anonymous, Environmental Protection Agency 2020). Many N-list decontaminants were listed as effective against SARS-CoV-2 with non-enveloped virus inactivation data but not SARS-CoV-2 data. This inclusion is because inactivation parameters for non-enveloped viruses were considered more challenging than enveloped viruses, hence decontaminants were accepted for SARS-CoV-2 if they had been successfully tested against non-enveloped virus. In the HHA work described here, the requirement for enveloped virus test results presents a similar objective to the EPA chemical decontaminant N-list insomuch as Φ6 was a greater challenge for HHA than the current virus of interest SARS-CoV-2. However, it is our opinion that enveloped virus test results in this case are more informative than non-enveloped virus test results since enveloped virus is the goal; the envelope is a critical virus structure and a likely inactivation target for HHA, and enveloped virus dilution/recovery/extraction methods are unique compared to non-enveloped viruses. Furthermore, HHA parameters for enveloped virus inactivation avoids the expense of sterilization, defined as spore inactivation, which requires substantially higher temperature and longer time.

There are numerous mathematical tools that can be used to analyze decontamination data. RSM models were very useful here since they provided simplified graphical data to rapidly adjust to changing goals for log reduction, time, temperature and humidity. This became very important since the HHA goals (requirements) were frequently re-defined by end users (aircraft maintenance authorities at Aircraft Mobility Command) over the past 20 years, and numerous goal changes were directed during the COVID19 pandemic. The original goals were to show enveloped virus inactivation of ≥7 log_10_ out of a ≥8 log_10_ challenge. This challenge level was set because measurements with high concentrations of microbes greatly increased the confidence in inactivation and it mitigated the risk of incomplete decontamination (Hamilton et al 2013). High challenge levels are important since exposure limits (infectious dosages) are not well defined for many viruses such as SARS-CoV-1 and SARS-CoV-2, and many decision makers who manage decontaminants/decontamination processes require the lowest risk possible. In addition, high virus titers have been commonly detected from clinical samples including 8 log_10_ of coronavirus from nasal swabs (Leung et al. 2020) further justifying the 8 log_10_ challenge. Although HHA does not fall under the EPA’s regulatory jurisdiction because it is not classified as a chemical disinfectant, the inactivation goal was reduced from 7-log_10_ to 3-log_10_ inactivation by the funding agencies during the COVID19 pandemic to match the EPA N-list for decontaminants, and provide decontaminant options to end users. The HHA RSM showed 90% probability levels of inactivation for log reductions of 0 through 7 log_10_, which allowed for rapid adjustments during the aircraft testing.

There were originally no time limitations set for aircraft sterilization (spore decontamination). The HHA time goal was reduced to ≤24 h for enveloped virus to increase HHA throughput and user acceptance. The time goal was further reduced by the end users and funding agencies to “1 h hold time; < 6 h total time” after the COVID19 pandemic had begun.

The HHA temperature goal was originally capped at the accepted temperature limit of C-130 aircraft (80°C), but that was for sterilization (spore decontamination). The HHA temperature goal was reduced to 60°C for enveloped virus to increase HHA acceptance over a range of aircraft since 60°C is a standard, accepted test temperature for most aircraft materials. The time goal of ≤24 h provided the rationale for measuring HHA over several hours to develop RSM models. However, the end users and funding agencies raised the accepted temperature limit to 68.3°C during COVID19 to reduce decontamination time.

Although there was a strong preference to reduce/eliminate humidity from HHA to reduce cost and complexity, all of the experimental data here showed that Φ6 was stable after drying. Virus was quickly inactivated after it was diluted in aqueous solutions and heated (solution control data). For HHA, the combination of high heat and high humidity, i.e. a high dew point, was critical to virus inactivation on materials in hour-long timeframes. This is consistent with long-known observations that enveloped virus is stabilized after drying on fomites. As an example, the enveloped virus smallpox was stabilized after it peeled/sloughed along with host cells, and then dried on porous materials including clothes and blankets. Furthermore, historic data shows that enveloped viruses were stabilized in cold, dry climates (Fenn, 2001).

The end-goal of the RSM was simplistic graphs where all decontamination test variables could be compared, understood and utilized for field testing and field application. This was particularly useful for large hot, humid air decontamination systems with fluctuating temperature and RH, potential power disruptions, and decontamination goals that have fluctuated widely over time. Since the RSM models showed little difference in inactivation kinetics among the different materials, a composite RSM was developed as a primary reference guide. This reference guide served the purpose of guiding HHA field-testing in aircraft. The RSM models translated to field demonstrations because there was a ≥5.9-log_10_ enveloped virus inactivation after both aircraft field tests, as predicted. The RSM models were also used as a guide for a number of iterative improvements between the first and second aircraft demonstration that shortened the overall HHA decontamination time from 5.5 to <3.5 h.

There were numerous conclusions. The Φ6 test indicators were useful measurements of inactivation for both lab and field-testing. HHA successfully inactivated enveloped virus at high temperature and high humidity. The RSM models were adaptable to changes in user requirements and successfully utilized to predict HHA results for field applications. Much methods development and testing remains to confirm HHA inactivation kinetics across virus species, and to develop HHA models that include HHA ramp-up and ramp-down times. Lastly, HHA decontamination of aircraft was successful because the enveloped virus on the C-17 aircraft was inactivated, the aircraft was flown after the HHA field test, and then the C-17 was certified as an operational aircraft.

## ACKNOWLEDGEMENTS

This work was supported through funding provided by the Defense Threat Reduction Agency (DTRA), Hazard Mitigation Capability Area (BA2 and 3 funds, Project Number CB10141) and Joint Program Executive Office CBRND/Joint Program Manager-CBRN Protection/Joint Biological Agent Decontamination System Program (JBADS). Thanks to Dr. Tabitha Newman for technical support. The aircraft demonstration events involved a consortium of supporters including DTRA, JPEO-CBRND, Aeroclave, LLC, Air Force, Air Mobility Command, United States Transportation Command, Naval Surface Warfare Center – Dahlgren Division, Air Force Research Lab, METSS Corporation, CCDC and Dover Air Force Base.

